# The effect of *Citrus reticulata* peel extract containing hesperidin on inhibition of SARS-CoV-2 infection based on pseudovirus entry assays

**DOI:** 10.1101/2024.05.03.592493

**Authors:** Endah Puji Septisetyani, Hayfa Salsabila Harsan, Dennaya Kumara, Pekik Wiji Prasetyaningrum, Komang Alit Paramitasari, Anisa Devi Cahyani, Khairul Anam, Ria Fajarwati Kastian, Adi Santoso, Muthi Ikawati, Edy Meiyanto

## Abstract

Orange (*Citrus reticulata* Blanco) peel contains a flavonoid glycoside hesperidin (HSD) as the primary component. HSD, upon enzymatic hydrolysis, forms hesperetin (HST) aglycone derivate. These two flavonoids have been predicted to have *in-silico* affinities for ACE2 and SARS-CoV-2 spike, crucial proteins in SARS-CoV-2 infection mechanisms. However, *in vitro* antiviral testing of orange peel extract, HSD, and HST has not been reported. This study presents for the first time a pseudovirus entry assay approach to test the anti-SARS-CoV-2 effect of HSD, HST, and orange peel extract prepared by hydrodynamic cavitation (HCV). We used a non-virulent pseudovirus model as an alternative to the original virus to target the entry point and enable research to be conducted outside the BSL-3 facility. Based on HPLC analysis, the test results showed that HCV contained HSD at about 4% w/w. Moreover, HSD 1 and 10 μM, HST 10 μM, and HCV 1 μg/ml showed inhibition of pseudovirus entry in 293/hACE2 cells with percentages inhibition 25.92, 37.40, 27.32, and 38.97 %, respectively. Despite HCV 1 μg/ml showing about 6 % lower inhibitory activity than HSD 1 μM in pseudovirus entry assay, it holds potential as a supplement or source of raw material for HSD as a SARS-CoV-2 antiviral.

## 1. INTRODUCTION

The coronavirus disease 2019 (COVID-19) has caused an outbreak worldwide [1]. Even though WHO has declared the end of the COVID-19 pandemic, marking the transition into an endemic disease [2], new cases of COVID-19 are still emerging, accompanied by new mutations found in the viral genome of the causative virus SARS-CoV-2 [3]. SARS-CoV-2 infections generally cause mild to moderate symptoms, for instance, cough, sore throat, fever, and myalgia [4]. However, although different major clinical manifestations occur across SARS-CoV-2 variants, more persistent symptoms are observed for current Omicron variants such as Omicron BA.2, which may affect long-term routine activities [5,6]. On the other hand, the main focus of the global response to this disease has been to minimize the risk of SARS-CoV-2 transmission by vaccine development [7]. Nevertheless, as SARS-CoV-2 has evolved, developing therapeutic candidates that can prevent COVID-19 progression becomes more crucial.

Citrus (*Citrus reticulata* Blanco) peel is rich in hesperidin (HSD) [8,9]. Upon enzymatic hydrolysis, HSD is converted to its aglycon, hesperetin (HST) (Figure 1A), which has a better membrane permeability and is more absorbable than HSD [10]. In-silico prediction of anti-SARS-CoV-2 effects of these flavonoids shown by HSD and HST binding affinities to ACE2 and TMPRSS2 [11-13], two proteins involved in SARS-CoV-2 cellular entry [14,15]. However, a report on laboratory-based *in vitro* assays showing HSD-HST antiviral effects is still lacking. On the other hand, since SARS-CoV-2 is considered a risk group 3 virus by the WHO [16], a biosafety level (BSL) 3 containment is needed to handle this virus. Therefore, the limited number of facilities providing BSL-3 containments becomes a challenge in studying SARS-CoV-2.

**Figure 1.**
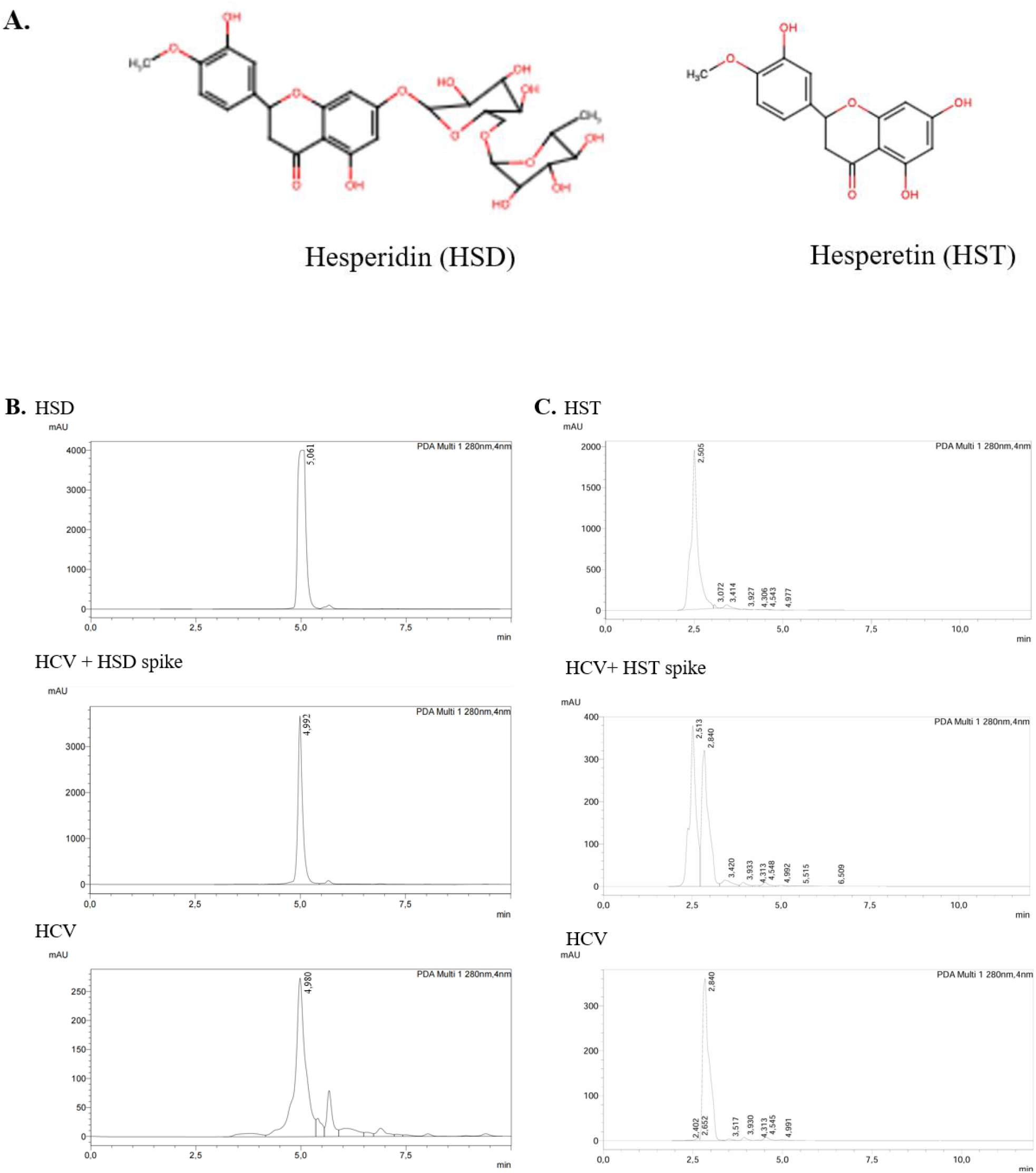
Hesperidin (HSD) and hesperetin (HST) contents in the orange peel (HCV) extract. A. Structure of HSD and HST. B. Representative HPLC chromatograms of standard HSD, HCV extract, and HSD spiked HCV extract. C. Representative chromatograms of standard HST, HCV extract, and HST spiked HCV extract.

The pseudovirus model offers an alternative to performing a SARS-CoV-2 cellular entry assay. Pseudovirus is a virus whose main structure and envelope proteins are derived from two viruses [17]. The main structure of SARS-CoV-2 pseudovirus can be constructed from a lower-risk group virus, such as vesicular stomatitis virus (VSV), enabling it to be handled in a laboratory with a lower biosafety level [18]. As a primary structure, VSV can be engineered to incorporate spike proteins of SARS-CoV-2 as its outer proteins [19]. Thus, the SARS-CoV-2 pseudovirus model facilitates the study of HSD and HST effects on SARS-CoV-2 infection.

To observe the inhibitory effect of citrus peel extract, HSD, and HST on SARS-CoV-2 infection, we performed a SARS-CoV-2 pseudovirus internalization assay in 293T cells expressing hACE2/TMPRSS2 and in 293/hACE cells. A replication-incompetent pseudovirus with spike*ΔG-GFP rVSV was used as a model of SARS-CoV-2 on the internalization assay.

## 2. METHOD

### 2.1. Preparation of orange peel extract, HSD, and HST

As previously described, orange peel extract was prepared by hydrodynamic cavitation of dried orange peel (HCV) [20]. The HCV extract was freeze-dried to obtain dry extract. HSD (cat no. PHR1794) and HST (cat no. SHBL8821) were acquired from Sigma (Sigma-Aldrich, St. Louis, USA). The HCV extract was dissolved in DMSO to prepare a 50 mg/ml stock solution for cell assay, while the flavonoid compounds were dissolved in DMSO to make a 50 mM stock solution. All solutions were stored at -20°C until used. Before cell treatment, the serial concentrations of samples were newly prepared in a culture medium with DMSO as cosolvent with a final concentration of less than 1%.

### 2.2. Cell Culture

The 293T (ECACC 12022001) and 293 (ECACC 85120602) cell lines were obtained from the National Research and Innovation Agency (BRIN, Indonesia). The 293T cells were maintained in High-Glucose Dulbecco’s-modified Eagle’s medium (Gibco, Billings USA) with supplementation of 10% FBS (Sigma-Aldrich, St. Louis USA) and antibiotics (100 μg/ml streptomycin and 100 U/ml penicillin). Recombinant 293 cells expressing hACE2 were generated by lentiviral vector and cultured in MEM medium (Sigma-Aldrich, St. Louis USA) with supplementation of 10% FBS, antibiotics (100 μg/ml streptomycin and 100 U/ml penicillin) (Invitrogen, Thermo), and 1% NEAA (Invitrogen, Thermo). The cell culture was stored in a humid incubator at 37 °C with 5% CO_2_. The introduction of recombinant plasmid DNA into cells was carried out by using a polyethyleneimine transfection reagent (PEI-Max, Polysciences).

### 2.3. HPLC analyses

Orange peel extract was dissolved in DMSO to a final concentration of 1,000 μg/ml, while HSD was dissolved in DMSO to a final concentration of 800 μM, and HST was dissolved in DMSO to a final concentration of 100 μM. The solution was filtered through a 0.22 μm nylon filter before HPLC analysis (modified from [21]). Three microliters of the sample were injected into an HPLC instrument (UFLC Shimadzu, Shimadzu Corp., Kyoto, Japan) with a C18 reversed-phase column (Shim-pack GIST C18 5μm 4.6×25 mm, Shimadzu, Shimadzu Corp., Kyoto, Japan) using methanol: acetonitrile: 0.1% acetic acid (89.3:10:0.7) as mobile phase with a flow rate of 1 ml/min at 35°C and detected using a photodiode array detector at 280 nm.

### 2.4. Recombinant plasmids

Plasmid pcDNA3.1-SARS2-Spike (Addgene #145032) [22] was used for pseudotyping to produce SARS-CoV-2 pseudovirus. We used pcDNA3.1-hACE (Addgene #145033) [22] and TMPRSS2 (Addgene #53887) [23] plasmids to engineer the target cells for the pseudovirus entry assay. Moreover, to generate recombinant 293/hACE2 cells, 293 cells were transduced with lentivirus produced utilizing pCMV-dR8.2-dvpr (Addgene #8455) [24]; pCAGGS-G-Kan (Kerafast EH1017) [25]; and (RRL.sin.cPPT.SFFV/Ace2.IRES-neo.WPRE(MT129) (Addgene#145840) [26] plasmids were co-transfected into 293T cells. The plasmids were propagated in *E. coli* and purified from the cell pellet using a plasmid maxi/midi-prep kit (Qiagen, Venlo, The Netherlands). The plasmid DNA concentration was determined using a micro-volume spectrophotometer (Thermo Fisher Scientific, Massachusetts, USA).

### 2.5. Immunofluorescence staining

About 3×10^4^ 293T cells per well of a 24-well plate were seeded on coverslips coated with 2% gelatin. The following day, cells were transfected with TMPRSS2 plasmids and incubated for about 18 hours. Cells were then washed with PBS and left overnight inside a CO_2_ incubator. For immunofluorescence staining, the cells were fixed with 4% paraformaldehyde for about 15 minutes, permeabilized with 0.2% Triton-X for 10 minutes, then blocked with 1% BSA/PBS for 30 minutes-1 hour at room temperature (RT). Then, the cells were incubated in an anti-TMPRSS2 antibody (Novus Biologicals NBP2-93322, USA) diluted in blocking buffer at 1:250 overnight at 4°C. After washing with PBS, the cells were incubated in an antirabbit secondary antibody conjugated with Alexa Fluor™-488 (1:1,000) (Abcam ab150077, USA) for 1 hour at RT. Next, the excess antibody was washed with PBS, followed by nuclei staining and cell mounting using a DAPI-containing mountant (Abcam Ab104139, USA). The fluorescence signal was observed using an Olympus-IX83 microscope (Olympus, Tokyo, Japan).

### 2.6. Western blot

The pellet of cells was lysed with a cold RIPA buffer (Abcam ab288006, USA) containing a protease inhibitor cocktail (Nacalai Tesque, Japan). The total protein concentration of the lysates was determined by BCA assay (Thermo Scientific, USA) to prepare about 10-40 μg protein mixed in Laemmli buffer. The proteins were resolved by SDS-PAGE and transferred onto the PVDF membrane. The membrane was incubated in 5% skim milk/TBS/0.05% tween-20 or 5% skim milk/PBS/0.05% tween-20 for 1 hour to minimize non-specific binding. Then, the membrane was incubated with primary antibody (anti-SARS-CoV-2 spike (Abcam, USA) 1:1,000, anti-hACE2 (Sigma, USA) 1:1,000, or anti-α-actin (Sigma, USA) 1:2,000) overnight at 4°C. After washing, the membrane was incubated in HRP-conjugated anti-rabbit or anti-mouse secondary antibodies (Abcam ab205718 or ab6728) or ALP-conjugated anti-rabbit secondary antibodies (Abcam ab6722), 1:2,000 or 1:4,000, for about 2 hours at RT or overnight at 4°C. Chemiluminescence substrates (Abcam, USA) were added to visualize antibody reactions and observed using a chemiluminescence imaging system (Uvitec Cambridge, UK), while 1-Step™ NBT/BCIP Substrate Solution (Thermo Scientific 34042) was added to visualize antibody reaction based on colorimetric development.

### 2.7. SARS-CoV-2 pseudotyping

About 1.5×10^6^ of 293T cells were seeded on a 100 mm dish and incubated overnight. The cells were then transfected with SARS-CoV-2 spike encoding DNA plasmid and incubated for about 18 hours. The next day, the cells were incubated with pseudotyped G*ΔG-GFP rVSV (Kerafast EH1024-PM) [25] at MOI ∼3 for 1 hour inside a CO_2_ incubator [25]. Then, the medium containing pseudotyped rVSV virus was replaced with a culture medium added with anti-VSV-G antibody (1:2,000) (Invitrogen PA1-30138, Thermo Fisher Scientific, USA) to neutralize the excess of G*ΔG-GFP rVSV, and the cells were kept inside an incubator overnight. After pseudotyping, the cell-conditioned medium (CM) was collected and spun to remove cell debris. The supernatant containing SARS-CoV-2 spike*ΔG-GFP rVSV (SARS-CoV-2 pseudovirus) was stored at -80°C in aliquots.

### 2.8. Pseudovirus entry assay

About 4×10^4^ of 293T cells/well were seeded onto a gelatin-coated 8-chamber slide (SPL Life Sciences, Pyeongtaek, South Korea) and incubated overnight. The next day, the cells were transfected with hACE2 and TMPRSS2 encoding plasmids and incubated for about 18 hours. The cells were pre-treated with HSD, HST, or orange peel extract for 30 minutes, then treated with a combination of tested sample and SARS-CoV-2 pseudovirus at 1:2 ratios in 300 μl total volume and incubated for another 16-18 hours. At the end of incubation, the cells were fixed using 4% PFA and mounted with a DAPI-containing mountant. Pseudovirus internalization, represented by observing GFP spots, was investigated using an Olympus IX83 motorized fluorescence microscope (Olympus, Tokyo, Japan). The GFP spots were counted from 8 images for each treatment by Fiji software (National Institute of Health). Pseudovirus entry assay was also performed utilizing 293/hACE2 stable cells generated using a lentiviral vector. The cells were seeded onto a gelatin-coated 8-chamber slide and incubated overnight. The following day, the cells were treated with tested samples described above. GFP spots were analyzed using cellSense dimension imaging software (Olympus, Tokyo, Japan).

### 2.9. Statistical analysis

Analyses were performed using GraphPad Prism software (GraphPad software, Inc., California, USA). Data were presented as mean ± SEM, as indicated in each figure. For pseudovirus entry assay in 293T/hACE2/TMPRSS2 cells, one-way ANOVA using Tukey’s test was used to determine the statistical differences with a p-value <0.05 considered significant. While for pseudovirus entry assay in 293/hACE cells, statistical difference was analyzed using one-tail unpaired t-tests with a p-value <0.05 considered significant.

## 3. RESULT AND DISCUSSION

### 3.1. Determination of HSD and HST content in the orange peel extract

The orange peel contains bioflavonoids, especially HSD, and a smaller amount of its hydrolysis product, HST. We determined the HSD and HST contents using the HPLC method in tangerine peel extract prepared using hydro cavitation-based extraction (HCV). Based on Figure 1B, standard HSD appeared with a retention time of 5.06 minutes, while HSD in orange peel extract appeared with a retention time of 4.98 minutes. To ensure that the main peak in the extract was HSD, we spiked HSD in the sample extract and obtained a retention time of 4.99 minutes. By the appearance of the single main peak, it has been confirmed that the main peak in the HCV extract is HSD. The slight differences in the retention time can be due to the diverse extract content. Then, based on the area under curve (AUC) data, we calculated the HSD content in the DMSO soluble fraction of the extract and obtained an HSD content of 4.34% w/w extract.

With the same HPLC condition, we observed the peak of standard HST appeared with a retention time of 2.51 minutes, while the main peak of the HCV extract existed with a retention time of 2.84 minutes. When we spiked the standard HST to be run with HCV extract, both prominent peaks appeared separately with consistent retention times (Figure 1C).

### 3.2. Expression of recombinant SARS-CoV-2 spike, hACE2, and TMPRSS2

We performed Western blot and immunofluorescence staining to clarify the ectopic expression of the SARS-CoV-2 spike, hACE2, and TMPRSS2 in 293T cells. Using the anti-S1 sub-unit of spike WT antibody, we detected the expression of spike WT by Western blot (Figure 2A). Moreover, we also observed the spike WT expression by immunofluorescence staining (Figure 2B). In addition, hACE2 expression was also detected by Western blot (Figure 2C), while TMPRSS2 expression was detected by immunostaining (Figure 2D). Spike expression was expected for the pseudotyping of rVSV. Meanwhile, hACE2 and TMPRSS2 expressions were expected to modify the target cells for pseudovirus entry assay. SARS-CoV-2 spike pseudotyping analysis was already reported previously [27].

**Figure 2.**
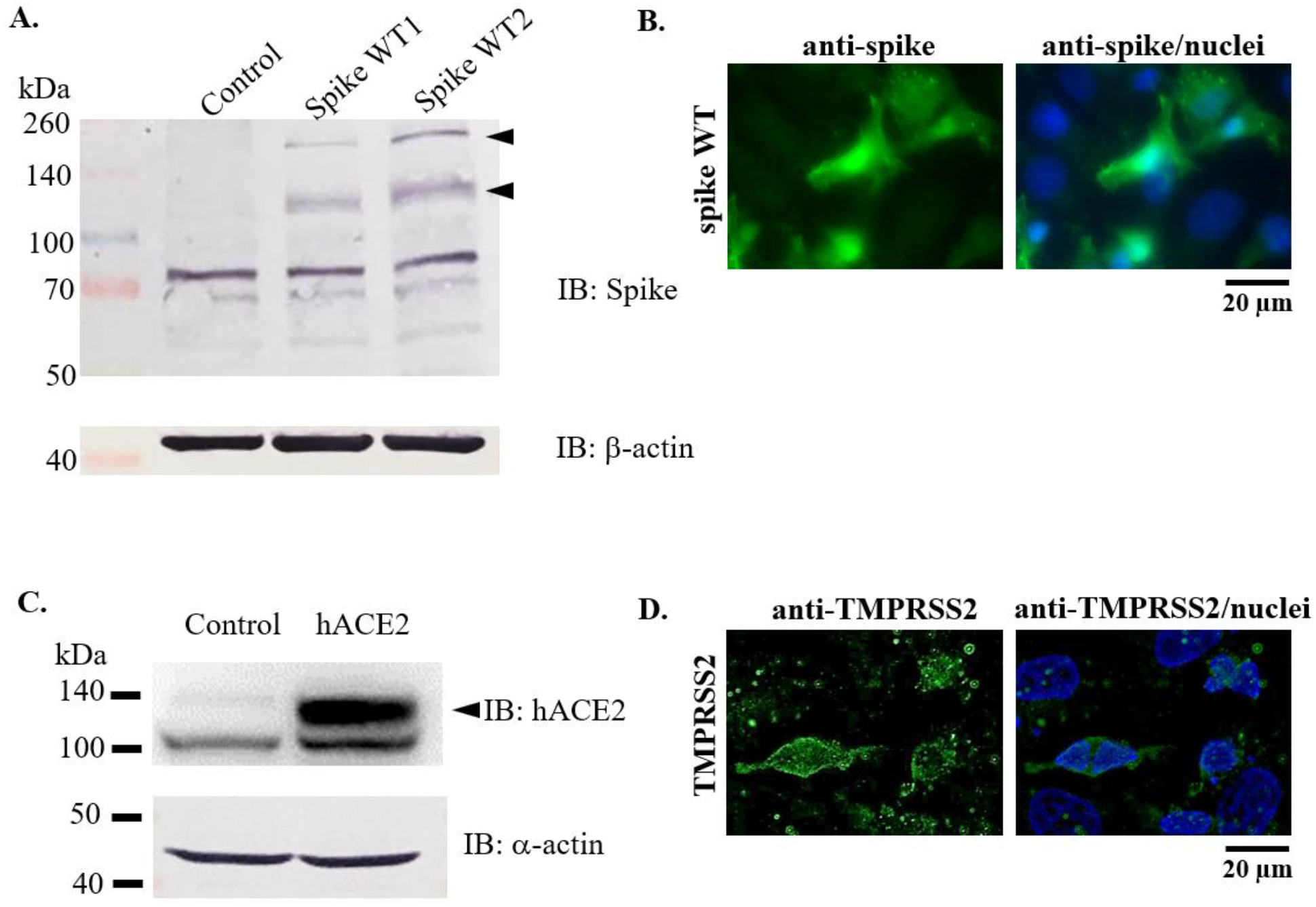
Confirmation of spike, hACE, and TMPRSS2 over-expression in 293T cells. A-C. Expression check of SARS-CoV-2 spike, hACE2, and β-actin by Western blot. D. TMPRSS2 detection by immunofluorescence staining.

### 3.3. Pseudovirus entry assay

The pseudovirus model was chosen to target the entry point of SARS-CoV-2. The pseudovirus also possesses spike peptide in its outer layer, which mediates binding to hACE2 as the target receptor, thus enabling the pseudovirus cellular entry. Moreover, the presence of TMPRSS2 will help pseudovirus internalization by priming the S1 subunit of the spike peptide. Then, upon internalization and release of the viral RNA genome, the GFP reporter will be expressed, generating a GFP spot. On the other hand, non-virulent pseudoviruses will not generate competent new pseudoviruses (Figure 3A). As shown in Figure 3B, the target cells were pretreated with a test solution before co-treatment with pseudovirus. Next, we investigated the GFP spots in the cellular area and analyzed the number of infected cells. As a result, the number of infected cells for control cells, HSD 10 and 100 μM, and HST 10 and 100 μM were 43.07, 14, 21, 18.25, and 15.83 %, respectively. Here, we showed that compared to control cells, HSD and HST treatment 10 and 100 μM can inhibit pseudovirus internalization at about 22-29 % (P<0.001) (Figure 3C).

**Figure 3.**
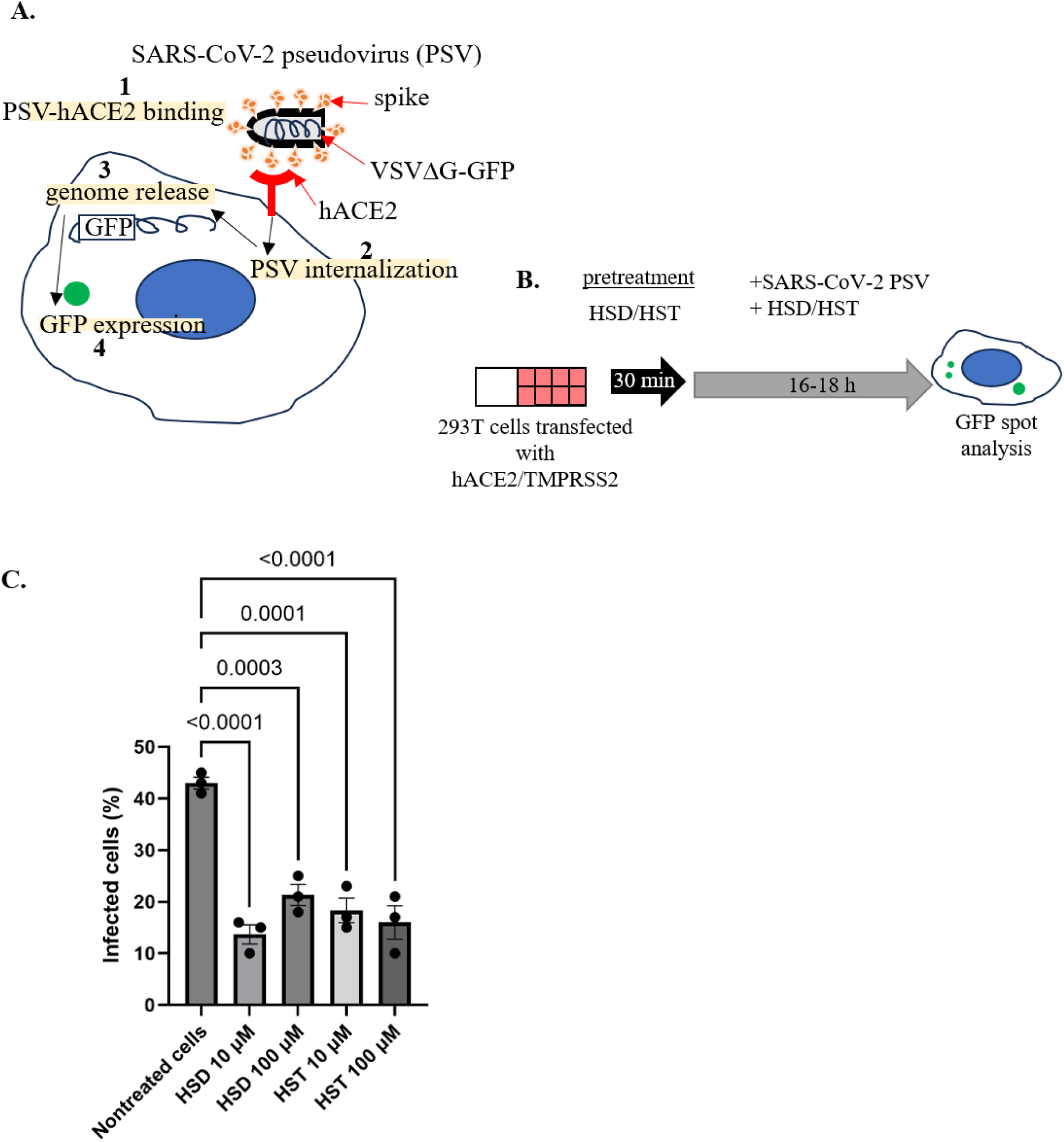
Effect of hesperidin, hesperetin, and orange peel extract on SARS-CoV-2 pseudovirus entry. A. Mechanism of SARS-CoV-2 pseudovirus cell entry and GFP expression. B. Schematic representation of pseudovirus entry assay. C. Quantification of SARS-CoV-2 (WT) pseudovirus entry assay in 293T cells transfected with hACE2/TMPRSS2. HSD = hesperidin, HST = hesperetin, PSV = pseudovirus. Data presented as mean ± SEM (n = 3 microscope fields).

Additionally, we performed a pseudovirus entry assay using 293/hACE2 stable cells. As in the previous experiment, cells were pretreated with the test material for 30 minutes, then treated with a mixture of pseudovirus and test material for 16-18 hours. Based on this assay, HSD demonstrated inhibition of pseudovirus entry at both 1 and 10 μM concentrations with the value of 34.28 % and 26.44 %. Meanwhile, HST showed the potential to inhibit pseudovirus entry at a concentration of 10 μM with a value of 30.69 % (P<0.05). When comparing the inhibitory effects of HSD and HST, HSD 1 μM and HST 1 μM show significantly different inhibitory effects on pseudovirus entry with 34.28% and 46.30% (P<0.05). HSD shows a more significant inhibitory potential than HST. Interestingly, HCV also showed inhibition, as did these two flavonoids. However, the inhibitory effect of 1 μg/ml extract was higher than that of 10 μg/ml extract concentration, with values 28.25 and 48.18 %, respectively (Figure 4).

**Figure 4.**
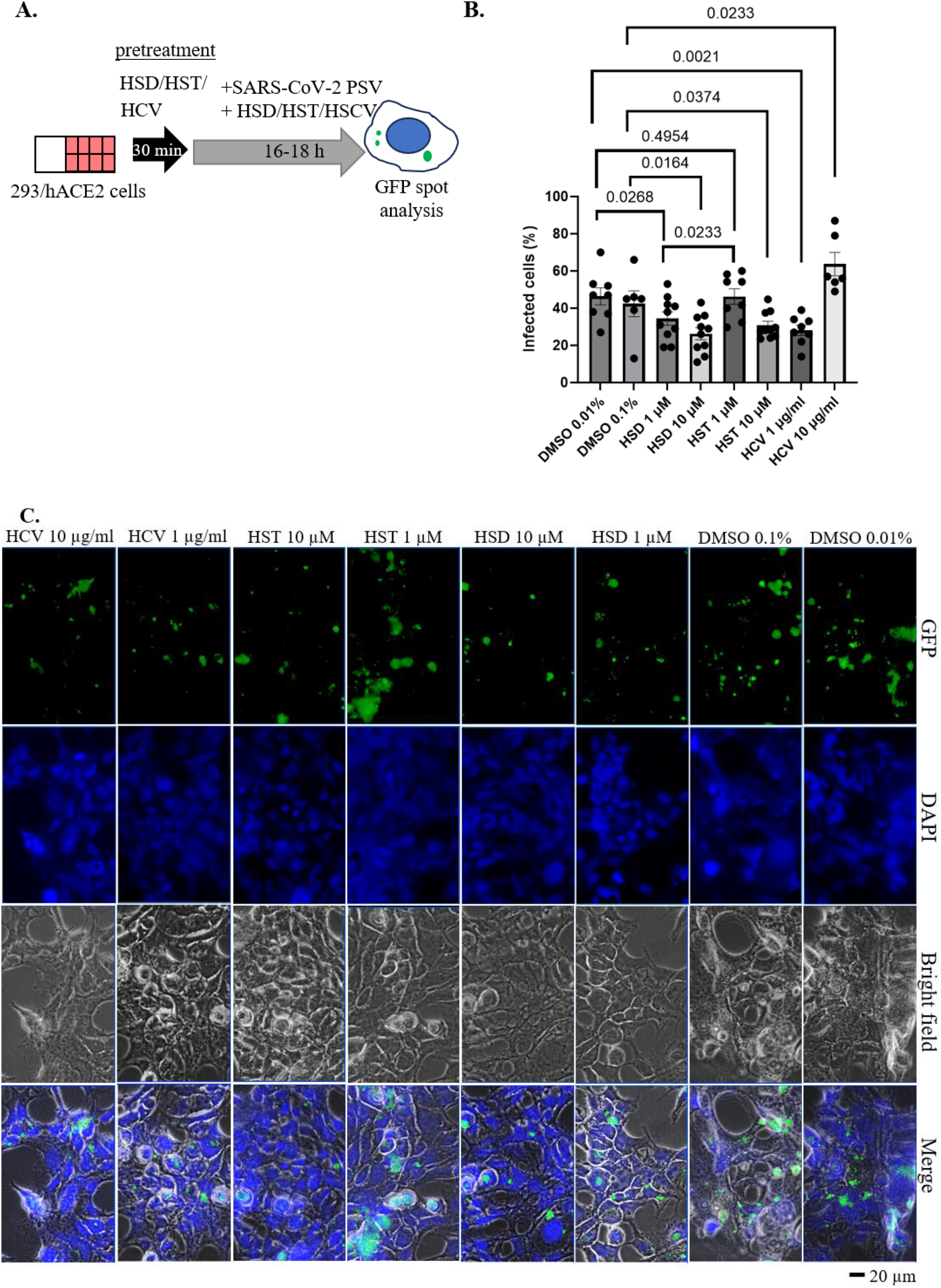
Effect of HSD, HST, and HCV on SARS-CoV-2 pseudovirus entry. A. Schematic representation of pseudovirus entry assay. B. Quantification of SARS-CoV-2 (WT) pseudovirus entry assay 293 cells stably expressing hACE2. C. Representative images showed GFP spots. HSD = hesperidin; HST = hesperetin; HCV = orange peel extract, PSV = pseudovirus. Data presented as mean ± SEM (n = 6-10 microscope fields).

HSD and HST show the binding potential to the hACE2 receptor through 3 binding sites with docking energies of -4.21 kcal/mol and -6.09 kcal/mol, respectively, based on in silico molecular docking studies [28]. In addition, other molecular docking studies reported the binding potential of HSD and HST to spike glycoprotein-RBD (PDB: 6LXT) and the RBD-hACE2/PD-ACE2 complex (PDB: 6VW1). When compared to previous docking studies, this in silico study shows that the binding potential of HSD is higher than HST, both in the RBD spike or the RBD-hACE2 complex with HSD docking scores of -9.61 and -9.50 kcal/mol, while the docking score of HST is -9.08 and -6.71 kcal/mol. Compared with other citrus flavonoids, the binding potential of HSD and HST at these two receptors is more remarkable than tangeretin, nobiletin, and naringenin [11].

Referring to the pseudovirus entry assay results on HSD and HST, these two compounds exhibit pseudovirus entry inhibitory activity in hACE2 expressing 293T cells with comparable effects without showing a statistically significant difference (P>0.05) (Figure 3). However, the inhibitory effect of pseudovirus entry on 293/hACE stable cells from HSD and HST appeared significantly different at a low concentration of 1 μM (Figure 4). In addition, HCV containing HSD and HST showed potential for pseudovirus inhibition. Apart from HSD and HST, standardized orange peel extract will be beneficial to reduce the risk of SARS-CoV-2 viral infection.

A non-competent pseudovirus is a non-virulent virus model that is very advantageous for studying the antiviral effect of targeting the viral entry point. The pseudovirus entry assay will represent the interaction between SARS-CoV-2 spike proteins incorporated in the outer part of the VSV backbone with hACE2 of the target cell [19]. This specific mechanism restricts the use of pseudovirus assay for studying viral replication. However, based on molecular docking studies, HSD and HST are predicted to interact with the SARS-CoV-2 protease (PDB ID: 6LU7), which is expected to inhibit virus replication [28, 29]. However, a lab-based assay is still needed to prove the inhibitory effect of those compounds on SARS-CoV-2 replication.

## 5. CONCLUSION

*C. reticulata* peel contains HSD as a major component [9]. We can also confirm the significant HSD content in the orange peel HCV extract we tested in this study. Since we could not confirm the HST content in the extract, the HSD content can be used to standardize the HCV extract for further research. In addition, besides being absorbed as HST, HSD can also be absorbed via the intestine without undergoing hydrolysis. This evidence was shown by detecting conjugated or non-conjugated hesperidin in the urine of rats fed with hesperidin [30].

Another flavonoid in orange peel, naringin, has been studied for its potential to prevent cytokine storms by decreasing the expression of LPS-induced proinflammatory cytokines COX-2, iNOS, IL-1β, and IL-6 [9]. Naringin also inhibits SARS-CoV-2 pseudovirus entry at 100 μM in hACE2-overexpressing 293T cells [27]. Apart from that, in the tests carried out in the pseudovirus entry, these orange peel HCV extracts inhibited SARS-CoV-2 infection. Therefore, it strengthens the potential for using orange peel extract to prevent SARS-CoV-2 infection or to reduce the severity of disease prognosis in COVID-19 patients.

## AUTHOR CONTRIBUTIONS

Conceptualization: EPS, EM; Data curation: EPS, HSH, DK, PWP, KAP, ADC. Writing - original draft: EPS, HSH, DK; Writing - review & editing: EPS, HSH, DK, MI, KA, RFK, AS, EM; Resources: EPS, EM.

## ACKNOWLEDGMENTS

We thank the iLab facility of the National Research and Innovation Agency (BRIN) for performing HPLC analyses.

## ETHICAL APPROVALS

Not applicable.

